# Dependence of fluorodeoxyglucose (FDG) uptake on cell cycle and dry mass: a single-cell study using a multi-modal radiography platform

**DOI:** 10.1101/668392

**Authors:** Yongjin Sung, Marc-Andre Tetrault, Kazue Takahashi, Jinsong Ouyang, Guillem Pratx, Marc Normandin, Georges El Fakhri

**Affiliations:** College of Engineering and Applied Science, University of Wisconsin, Milwaukee, WI 53211, USA; Gordon Center for Medical Imaging, Department of Radiology, Massachusetts General Hospital, Boston, MA 02114, USA; Department of Radiology, Harvard Medical School, Boston, MA 02115, USA; Department of Radiation Oncology and Medical Physics, Stanford University, Stanford, CA 94305, USA

## Abstract

High glucose uptake by cancer compared to normal tissues has long been utilized in fluorodeoxyglucose-based positron emission tomography (FDG-PET) as a contrast mechanism. The FDG uptake rate has been further related to the proliferative potential of cancer, specifically the proliferation index (PI) − the proportion of cells in S, G2 or M phases. The underlying hypothesis was that the cells preparing for cell division would consume more energy and metabolites as building blocks for biosynthesis. Despite the wide clinical use, mixed reports exist in the literature on the relationship between FDG uptake and PI. This may be due to the large variation in cancer types or methods adopted for the measurements. Of note, the existing methods can only measure the average properties of a tumor mass or cell population with highly-heterogeneous constituents. In this study, we have built a multi-modal live-cell radiography system and measured the [^18^F]FDG uptake by single HeLa cells together with their dry mass and cell cycle phase. The results show that HeLa cells take up twice more [^18^F]FDG in S, G2 or M phases than in G1 phase, which confirms the association between FDG uptake and PI at a single-cell level. Importantly, we show that [^18^F]FDG uptake and cell dry mass have a positive correlation in HeLa cells, which suggests that high [^18^F]FDG uptake in S, G2 or M phases can be largely attributed to increased dry mass, rather than the activities preparing for cell division. This interpretation is consistent with recent observations that the energy required for the preparation of cell division is much smaller than that for maintaining house-keeping proteins.

## Introduction

Most biological systems including humans rely on glucose as a source of energy. Cancerous cells tend to take up more glucose than normal tissues to support their fast growth while relying on an inefficient route, fermentation instead of respiration, to generate energy^1^. This higher glucose uptake by cancer compared to normal tissues has been utilized in positron emission tomography (PET)^2^. [^18^F] fluorodeoxyglucose ([^18^F]FDG), a glucose analog labeled with fluorine-18, can be delivered into cells by the same transporter system used by glucose, but, once inside the cell, it is phosphorylated to a form that prevents further metabolism and release from the cell^3^, and thus its accumulation can serve as an imaging contrast. Fluorine-18 has a 110 min half-life and decays by positron emission 97% of the time. The positrons emitted from this radionuclide have maximum and mean kinetic energies of 635 and 250 keV, respectively; however, once their energy is reduced to a few eV due to scattering, the positrons annihilate with electrons to generate pairs of 511 keV gamma rays. PET detects these highly energetic gamma rays using an array of scintillator detectors and, after tomographic reconstruction, provide the distribution of [^18^F]FDG within the body, and thus the locations of metabolically-active tissues such as cancer in three dimensions^4^. Of note, the annihilation gamma rays are generated 0.4–2.0 mm away from the radionuclides—a positron range that fundamentally limits the spatial resolution of PET^5^.

The most widespread application of PET has been for the detection and staging of cancer^5^. Standardized uptake value (SUV), a measure of FDG uptake, has been related to the proliferation index (PI), a measure of the proliferative potential of cancer^6^. Specifically, PI is defined as the proportion of the cells in S, G2 or M phases of the cell cycle^7,8^. Despite the success of PET as a cancer diagnostic tool and its ever-increasing use for prognosis, the relationship between PI and FDG uptake (or SUV) is not well established. This may be because PET measures the average activity within a tissue region of interest, while a tumor has a highly heterogeneous distribution of cancer cells with the morphological and physiological features affected by the local microenvironments^9^. The cells in a population can also show highly heterogeneous responses to external stimuli depending on various parameters.

The present paper establishes a multi-modal radiography platform to study the relationship between FDG uptake and various cell parameters at a single-cell level. Using our new system, we have measured [^18^F]FDG uptake by single HeLa cells together with their dry mass and cell cycle phase. The dry mass of a cell is the result of growth and metabolism, and thus we hypothesize that it is related to the rate of glucose uptake. The measurement of cell cycle phase allows us to study the relationship between [^18^F]FDG uptake and PI at a single-cell level.

## Materials and Methods

### Cell culture

HeLa human cervical cancer cells transfected in conjunction with fluorescence ubiquitination cell cycle indicator (FUCCI) cell cycle sensor (Life Technologies, P36237) were kindly donated by Dr. Seungeun Oh (Department of Systems Biology, Harvard Medical School). The cells stably express two cell cycle-regulated proteins, geminin and Cdt1, fused to one green (emGFP) and red (TagRFP) fluorescent protein. The cells were cultured in Dulbecco Modified Eagle medium (Invitrogen, 21063-029) supplemented with 10 % FBS (Invitrogen, 10438026) and 1 % 100x penicillin-streptomycin solution (Invitrogen, 15140-122).

### Multi-modal live-cell radiography system

We have built a multi-modal live-cell imaging platform based on an upright fluorescence microscope (Nikon E800, Figure 1). It supports sequential acquisitions of the same field of view using bright field imaging, fluorescence imaging, quantitative phase imaging and radioluminescence imaging. They respectively provide information on cell morphology, cell cycle stage (by utilizing FUCCI cells), cell dry mass and quantitative radionuclide uptake.

**Figure 1.**
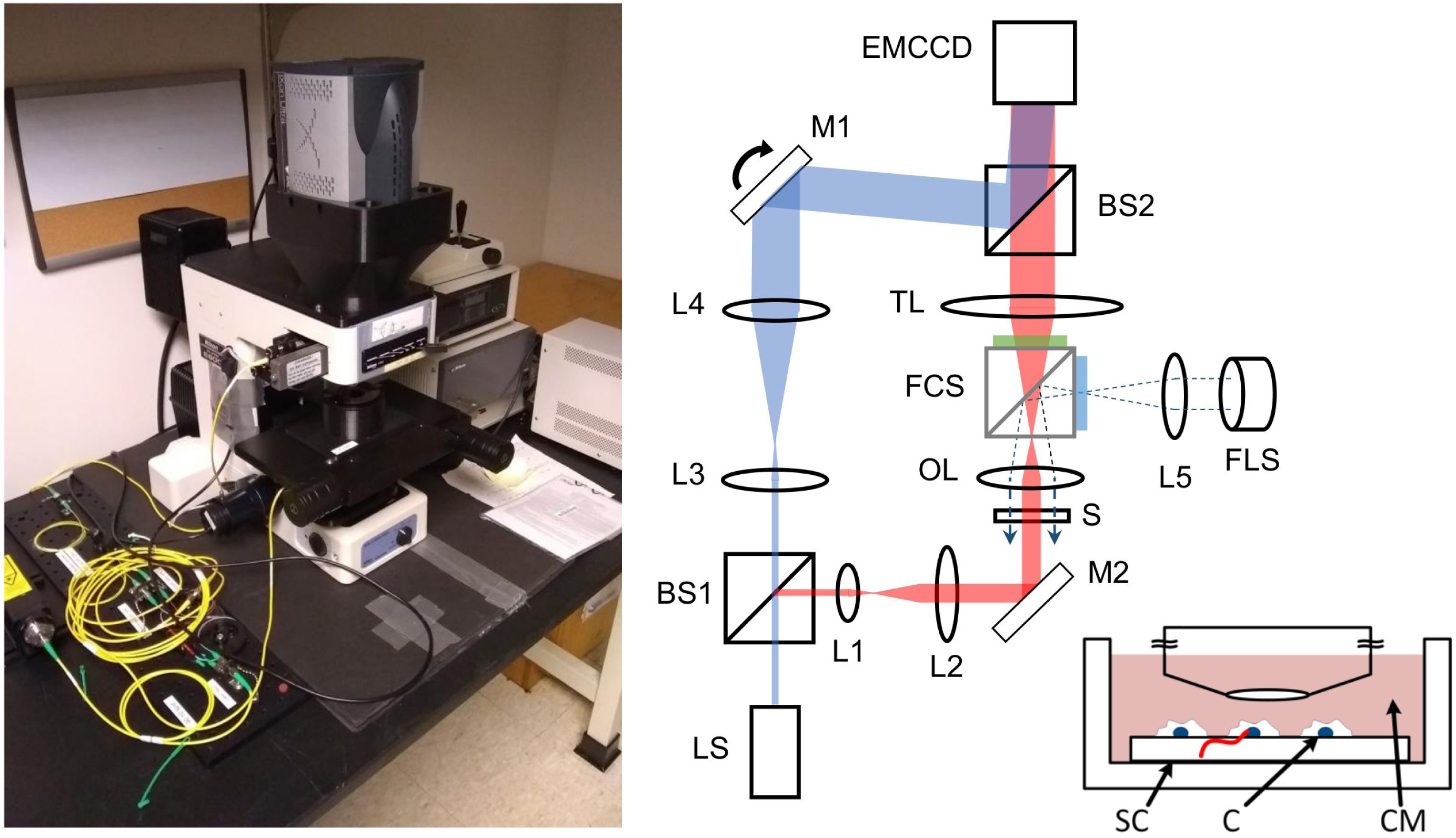
Multi-modal live-cell radiography system and example images. LS: light source (laser); FLS: fluorescence light source (Mercury lamp) BS1, BS2: beam splitter; L1-L5: lenses; M1, M2: mirrors, S: sample; OL: objective lens; TL: tube lens; FCS: fluorescence filter cube slider; EMCCD: EM-CCD camera. The FCS is inserted into the beam path only for fluorescence imaging. The dotted lines represent the excitation beam path. The inset shows a radiolabeled cell (C) grown on a scintillator crystal (SC), which is immersed in culture medium (CM) during measurement. The red line represents the ionization track in the crystal.

The platform relies on a high-magnification imaging chain (x40, 0.8-NA objective lens coupled to 105-mm tube lens) and an electron-multiplying-charge-coupled-device (EMCCD) camera (Andor, iXon Ultra 888) with high pixel resolution and sensitivity. At full resolution and this magnification level, the camera’s 1024 × 1024 sensor array obtains a resolution of 0.625 µm per pixel, for an overall field of view of 640 µm by 640 µm. The camera is the common image recording device, although specific settings are tailored for each imaging modality.

Bright field imaging relies on a standard upright microscope configuration illuminated by a halogen lamp and provides cell morphology information. No special configuration was introduced for this imaging modality. A standard fluorescence configuration based on a metal-halide arc lamp was used to obtain cell cycle information. The system determines the cell cycle phase by two-channel fluorescence imaging of FUCCI reporters. The green fluorescent protein (GFP) of the FUCCI reporter has its excitation peak at 488 nm and its emission peak at 510 nm, whereas the red fluorescent protein (RFP) has its excitation peak at 555 nm and its emission peak at 584 nm. For the GFP measurement, we used a fluorescence filter set (Nikon, B-2E) with an excitation band of 450 – 490 nm, an emission band of 520 – 560 nm, and a cut-on wavelength for the dichroic mirror at 505 nm. For the RFP measurement, we used an excitation filter (Thorlabs, MF559-34) with a center wavelength (CWL) of 559 nm and a bandwidth of (BW) 34 nm, an emission filter (Thorlabs, MF620-52) with a CWL of 620 nm and a BW of 52 nm, and a dichroic mirror with a cut-on wavelength of 596 nm.

Quantitative phase microscopy (QPM) was used to measure a 2-D map of optical phase alteration induced by individual cells. QPM is a wavefront-sensing technique adapted to imaging microscopic specimens. The wavefront distortion or the phase alteration is proportional to the refractive index, which is related to the dry density, the concentration of non-aqueous contents within the cell. QPM can be built using an interferometer^10–12^, a Shack-Hartmann sensor^13,14^, or a transport-of-intensity method^15,16^. In this study, we used an off-axis interferometer (Figure 1), which allows acquiring the phase image in a single snapshot. A beam splitter (BS1) was used to split a collimated beam into two: sample and reference beams. To incorporate the QPM into the microscope body, we split the beam using a 2 × 2 wideband fiber optical coupler (Thorlabs, TW630R5A2). The sample and the reference beam used in QPM are shown in red and blue, respectively, in Fig. 1. The second beam splitter (BS2), which combined the sample and the reference beams, was mounted on the fluorescence filter cube slider. The reference beam was slightly tilted with respect to the sample beam, which generated straight fringes on the detector when there was no sample. The fringe pattern was distorted when the phase distribution in one beam was altered with respect to that in the other beam. Thus, using the reference beam, which had a uniform phase distribution, we measured the sample-induced phase alteration from the recorded fringe pattern. Figure 2(a) shows an example raw interferogram acquired with QPM. The small, rectangular region in the figure is enlarged on the right, which clearly shows straight fringes in the background region and their distortion near the cell. A more complete and detailed description on QPM can be found in our previous publications^17,18^.

**Figure 2.**
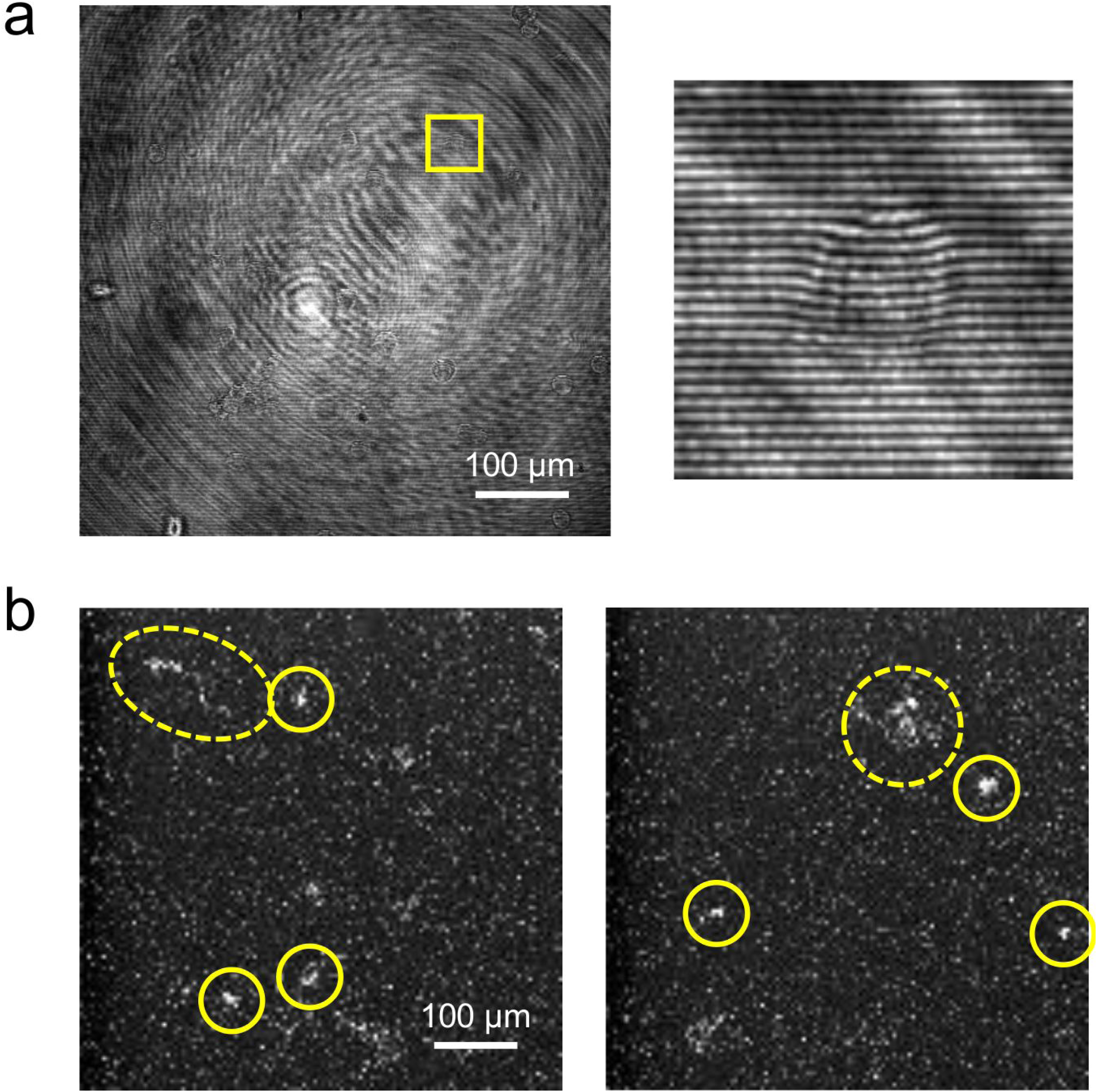
Example raw images acquired with quantitative phase microscopy (QPM) and radioluminescence microscopy (RLM). (a) An example QPM image with the rectangular region enlarged on the right. (b) Example RLM images, where short ionization tracks are encircled with solid lines and long or diffuse ionization tracks encircled with dotted lines.

Lastly, radioluminescence microscopy (RLM) is a low-light imaging modality^19,20^. Its operating principle resides on acquiring the light generated by beta ionization traveling through a scintillating crystal in contact with the radio-labeled bio-sample. By locating the tracks and counting them over a pre-determined acquisition time, the system can measure the 2D distribution of the tracer, analogous to PET or single-photon emission computed tomography (SPECT) molecular images. Since the system directly detects positrons rather than the resulting annihilation gamma pair, it is possible to reach spatial resolutions below the positron’s range, required for single-cell imaging. Of note, RLM typically uses a tube lens with short focal length (e.g., 50 mm) to increase the image brightness^21^. In the present setup, we used a tube lens with slightly longer focal length (105 mm) to incorporate the fluorescence filter cube slider at the cost of reduced sensitivity.

A dense scintillator is required to provide high stopping power and thus high sensitivity. The thickness of the scintillator determines both the spatial resolution and the sensitivity of RLM; thin scintillators yield finer spatial resolution, while thick crystals increase sensitivity. An affordable compromise was reached using CdWO_4_ crystals (MTI Corp., CWO4g050501S2) with dimension of 5 × 5 × 0.1 mm (length × width × thickness), polished on both sides by the manufacturer. Previous simulations and experiments using these crystals obtain spatial resolutions of approximately 30 µm^22^. Knowing this, the camera can be used with maximum binning (8 × 8), resulting in 5 µm per pixel image resolution. This change in settings maximized the camera’s signal to noise ratio without degrading the spatial resolution, a strong benefit in RLM’s extremely low light conditions. Figure 2(b) shows example raw images acquired with RLM. The figure shows short ionization tracks (solid lines) as well as a long or a diffuse ionization track (dotted lines). Short ionization tracks are produced by positrons traveling perpendicular to the imaging plane; thus, the center of mass can be approximated to the location of the radionuclide that produced the positrons. Long ionization tracks are produced by positrons incident onto the crystal surface at a shallow angle. We discarded them in favor of shorter tracks that are easier to localize. Diffuse tracks are formed by gamma rays annihilated deep in the crystal, and thus were excluded from further processing^19^.

### Sample preparation and imaging procedure

HeLa-FUCCI cells were harvested at 70 - 80 % confluency in 25-cm^2^ culture flasks, then collected with 0.05 % Trypsin-EDTA (Thermo Fisher Scientific, 25300054), diluted to 1:100 in fresh culture medium, and plated onto the CdWO_4_ crystal in a glass-bottom dish (MatTek Corp., P35G-1.0-14-C.S). The crystal and dish were sterilized with a mixture of 70:30 ethanol/water (vol/vol), then washed three times with sterile distilled water, and then placed under ultraviolet light in a biosafety cabinet for an hour. The cells were incubated overnight in a 5 % CO_2_ cell culture incubator at 37 °C.

[^18^F]FDG produced by the cyclotron at the Massachusetts General Hospital (MGH) Gordon PET Core was mixed with glucose-free DMEM (Thermo Fisher Scientific, 11966-025) to the concentration of 2 mCi per mL at the beginning of experiment. The cells were placed for one hour in glucose-free DMEM to increase glucose consumption. Then, we aspirated the glucose-free DMEM from the dish, added 0.1 mL of the [^18^F]FDG solution to the well inside the dish, and returned it to the cell incubator. After another hour, we aspirated the hot solution, washed the entire culture area three times with warm DMEM to remove residual [^18^F]FDG, and imaged the samples with the multi-modal microscope as described below.

Due to the low spatial resolution, RLM requires the cells to be sparsely distributed. To optimize the number of cells imaged, we used bright-field imaging and manually searched for a field of view (FOV) containing many isolated single cells. After selecting a FOV and centering the focal plane on the cells, snapshots were sequentially taken using bright-field imaging, green and red fluorescence imaging, and QPM imaging, in this order (BF, GFP, RFP, QPM), with the Micromanager acquisition software^23,24^. The images were taken at full camera resolution, with exposure and gain adjusted by modality (see table I) to ensure good image dynamic range in each case. RLM acquisitions on the other hand record individual ionization tracks *in the crystal* over several snapshots. Therefore, the focal plane was positioned a few microns below the crystal surface nearest to the cells (estimated using BF imaging). After ensuring total darkness in the room, 20,000 images were recorded continuously with an exposure time of 20 ms and an maximum electron-multiplying (EM) gain of 1000. This took about 10 minutes, including approximately 32% dead time. This will be reduced to 8% in future studies thanks to a special frame-buffer mode in the camera.

**Table I.** Camera settings for the image acquisition of all imaging modalities.

|  | BF | GFP | RFP | QPM | RLM |
| --- | --- | --- | --- | --- | --- |
| Exposure time (ms) | 10 | 100 | 100 | 10 | 20 |
| EM gain | 3* | 200 | 200 | 3* | 1000 |
| Camera binning | 1 × 1 | 1 × 1 | 1 × 1 | 1 × 1 | 8 × 8 |
| Image count | 1 | 1 | 1 | 1 | 20,000 |
| Acquisition time (minutes) | - | - | - | - | 10 |

\* Minimum EM Gain

All the measurements were completed within 90 minutes of adding the [^18^F]FDG solution to the dish and within 30 minutes of washing the sample with warm DMEM. Background single frames were acquired for an empty FOV using all the imaging modalities and the same settings, except for RLM where 1000 frames were recorded instead of only one.

### Image processing

Bright field images were used for visual inspection, focal plane positioning, manual cell validation and general quality control, and therefore did not receive specific pre-processing or correction procedures.

The green and the red fluorescence images were first normalized with their own background images to compensate for nonuniform distribution of excitation light. For each image, the average count in an empty region was subtracted, and the entire image was thresholded to further suppress residual noise. Using FUCCI, the cell cycle for each cell was determined by the green and the red fluorescence intensities summed over each cell area. Cells expressing only the red fluorescence protein are in G1 stage; those expressing only green are in S, G2 or M stage; and those expressing both are in G1/S stage. The remaining cells are in M/G1. The FUCCI sensor did not identify any cells in M/G1 among the data we collected. This is partly because cells that have just divided tend to stay clustered together and we did not image such cells, due to the limited spatial resolution of RLM. We manually verified the cell cycle assignment by visual inspection of each fluorescence image and found that only one cell out of 26 was falsely assigned to G1/S due to background noise in the red fluorescence image. We manually changed the assignment for the cell to S, G2 or M.

The QPM raw interferograms were processed as described in our previous publication^18^, which produced a phase image as shown in Fig. 3(c). The dry mass *m* of a cell was calculated from the measured phase image Φ(*x, y*) using the following formula:

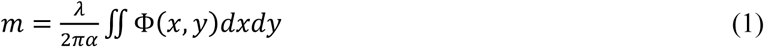

where *λ* is the laser’s wavelength, 633 nm for He-Ne laser, *α* = 0.18 mL/g is the specific refraction increment, and the integral is over the cell area, or cell boundary. We extracted this boundary by applying the Sobel edge detection method^25^ to the binary mask generated from the phase image. The phase image provided a clearer boundary than the bright-field image when the cell was flat.

**Figure 3.**
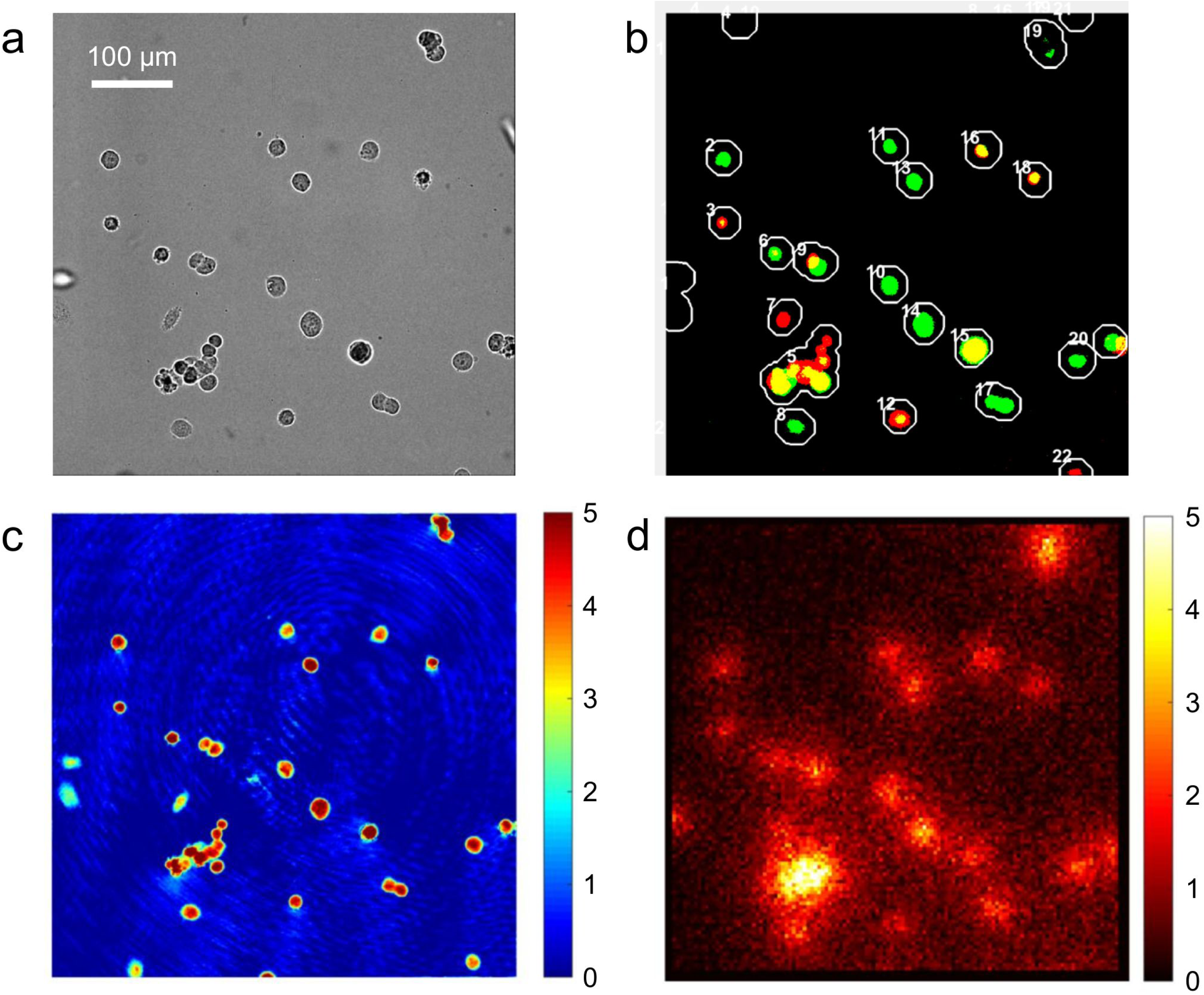
Example multi-modal images after image processing. (a) Bright-field image, (b) fluorescence image, (c) quantitative phase image (unit: radian), and (d) radioluminescence image of a HeLa cell (unit: counts/min). The fluorescence image in (b) was synthesized from an RFP and a GFP image, colored accordingly. In the image, the yellow color represents the expression of both the proteins, and the number next to each circle represents the cell label assignment used for visual inspection of the cell cycle.

After dark-field correction, radioluminescence images were processed using the ORBIT algorithm in MATLAB (The MathWorks, Inc.), downloaded from the link in Kim, et al.^21^. In individual frames, the algorithm identifies peaks corresponding to single positron scintillation events, calculates their center of gravity and increments a counter at that position. It also removes undesired events such as rare gamma events directly ionizing the EMCCD sensor, long tracks and gamma annihilation particles interacting in the crystal. The parameters used with the ORBIT are shown in Table 2. All 20,000 frames in an acquisition were processed to produce the radiotracer distribution image (Fig. 3d).

**Table II.**
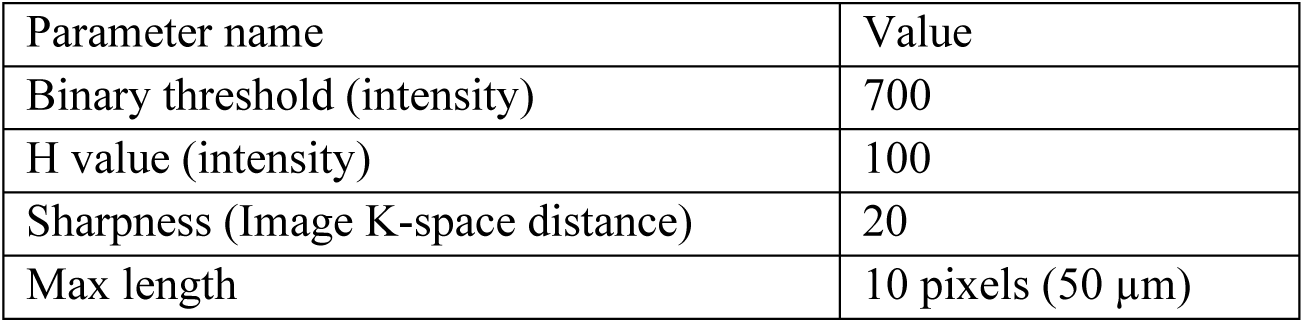
ORBIT reconstruction parameters.

Individual cell counts were obtained by integrating the RLM image values inside the cell boundary determined with the phase image (scaled to RLM’s 8 × 8 binning) but dilated to account for RLM’s spatial resolution. This dilation extends the boundary to encompass events generated by a cell but detected outside its volume. Furthermore, when this extension merges two cell boundaries, then the cells are considered too close because their counts potentially cause cross-contamination among each other. For our crystal dimensions, the spatial resolution is expected to be about 30 um FWHM^22^, or about σ = 12.5 um assuming a Gaussian distribution for the point-spread function. We selected a one sigma dilation radius to balance between gathering good counting statistics and keeping cells separated. The cells merged by the dilation were discarded from the experimental data (for example, boundary number 17 in Fig. 3b). The data were corrected for decay due to the time elapsed between the FDG’s calibration time and effective acquisition start in each experiment.

### Statistical Analysis

All measurements were expressed as interquartile range. Correlation of different cell cycle groups was computed using a two-tailed Mann–Whitney–Wilcoxon test and an ANOVA test. A P value of less than 0.05 was considered significant. For linear regression analyses, we used a robust regression which down-weights outliers according to the distance from the best-fit line and iteratively re-fits the model. Specifically, an iterated reweighted least squares (IRLS) algorithm was used with the Huber weights^26^. Statistical analysis and graph plotting were done using R (version 3.4.2).

## Results and Discussion

### Multi-modal images

Figures 3(a) through 3(d) show example images acquired with the multi-modal live-cell radiography system developed in this study. The bright-field image in Fig. 3(a) shows cell morphology. Figure 3(b) is a fluorescence image synthesized from an RFP and a GFP image. The cells colored in red express Cdt1 and thus are in G1 stage; the cells colored in green express geminin and thus are in S, G2 or M stage; the cells colored in yellow express both Cdt1 and geminin, and thus are in G1/S stage^27^. The cells not expressing either reporter are in M/G1. The number next to each circle represents the cell label identified by an automatic image segmentation algorithm in MATLAB. The phase image in Fig. 3(c) was used to calculate the cell dry mass as well as to generate a mask for the image segmentation. Figure 3(d) is a radioluminescence image synthesized from a 10-minute acquisition. The intensity represents the number of beta decays observed during the acquisition and is expressed in average counts per minute. Of note, the clustered cells in the left lower region are not distinguished in Fig. 3(d) due to the low spatial resolution of RLM.

### HeLa cells’ [^18^F]FDG uptake, cell dry mass, and cell cycle phase

The total number of HeLa cells considered for this study and observed in three separate experiments was 26. The median dry mass of the cells was 604 pg (interquartile range: 494−669), and the median number of counts per minute was 67 (interquartile range: 54−88). As shown in Fig. 4(a), there exists a linear and statistically significant relationship between [^18^F]FDG uptake and cell dry mass (adjusted R-squared = 0.46, *P* < 10^-4^). Figure 4(b) further shows that the mass-normalized uptake is not correlated with cell dry mass (adjusted R-squared = −0.04, *P* = 0.77). The total amount of [^18^F]FDG within a cell is the product of [^18^F]FDG concentration in the cell and cell volume. Assuming a constant cell density, the linear relationship between the [^18^F]FDG uptake and the cell dry mass suggests that intracellular FDG concentration is approximately the same.

**Figure 4.**
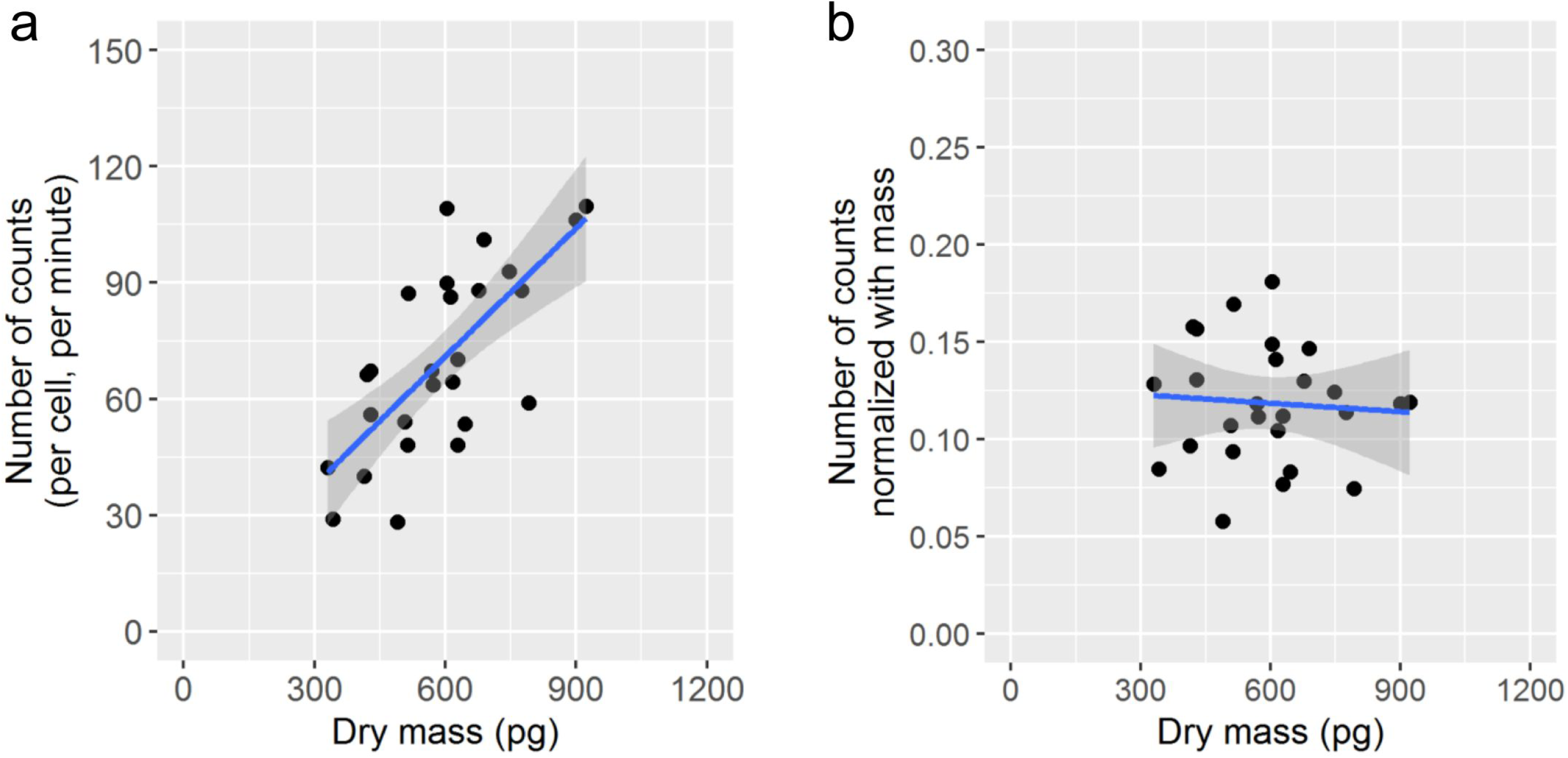
Linear relationship between [^18^F]FDG uptake and cell dry mass. (a) Scatter plot of the number of counts (per cell, per minute) vs. the cell dry mass, and (b) the same plot with the number of counts normalized with the cell dry mass. The number of counts is proportional to the [^18^F]FDG uptake. Regression lines are shown together with the 95 % confidence interval in gray.

Figures 5(a) shows the dry mass of the cells in different cell cycle phases, with Table III reporting statistically relevant information. The cells in S, G2 or M phases had significantly higher dry mass than the cells in G1 phase (*P* = 0.012), which is natural because the cells at a later stage (S, G2 or M) have a greater dry mass than the G1 cells. Figure 5(b) shows [^18^F]FDG uptake for cells in different cell cycle phases, and Table III includes related statistical analyses. Cells in the S, G2 or M phase had 97% higher [^18^F]FDG uptake than those in the G1 phase (*P* < 0.006). The number of counts divided by the cell dry mass did not show significant dependence on the cell cycle phase, as seen in Fig. 5(c).

**Table III.** Median and interquartile range for dry mass and [^18^F]FDG counts.

|  |  | All cells | G1 | G1/S | S/G2/M |
| --- | --- | --- | --- | --- | --- |
| Cell dry mass (pg) | Median | 604 | 422 | 612 | 628 |
|  | Interquartile range | 494–669 | 396-450 | 530-631 | 544-732 |
| [$^{18}\text{F}$]FDG decay counts (counts/min) | Median | 67 | 44 | 64 | 87 |
|  | Interquartile range | 54–88 | 37-50 | 48-77 | 65-95 |

**Figure 5.**
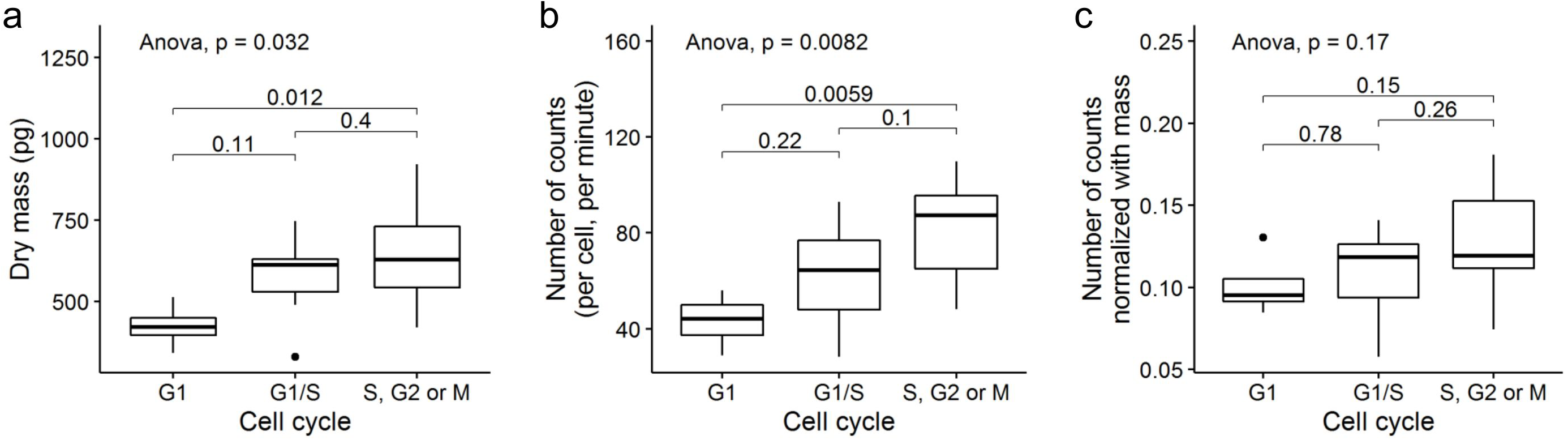
Cell cycle dependence of the dry mass (a), the number of counts (b) and the number of counts divided by the dry mass (c). Horizontal bars, boxes and whiskers represent the median, interquartile range and range respectively. P values are included for the analysis of variance (ANOVA) and two-tailed Mann-Whitney-Wilcoxon test.

The high glucose uptake in S phase by cells has been attributed to the high demand for nutrients and energy for synthesizing deoxynucleotides, DNA and histones^28^. For example, the uptake of 2-deoxy-D-[^3^H]glucose ([^3^H]DG) by transformed lymphocytes (IM9) is more than doubled during the S phase but, in G2 phase, it returns to the basal level of G1 phase^28^. The authors postulated that high [^3^H]DG uptake in IM9 cells was due to high demand for nutrients for synthesizing deoxynucleotides, DNA and histones - the materials required for cell replication^28^. However, recent studies demonstrated that the energy required for the synthesis of new materials in preparing for replication is much smaller than the energy required for normal cell operation^29,30^. It was also shown that carbon mass in cells is mostly derived from other amino acids rather than glucose^31^. Considering these observations, it is not clear why the cells in S phase take up twice more glucose than the cells in gap phases. On the other hand, the linearity between cells’ growth rate and their size is well established^32^. In our previous study, we showed that the growth rate increases almost linearly until the dry mass reaches a critical size^33^. Considering that cellular growth is an energy-demanding process and glucose is a major source of energy, it seems evident that the large cells in the later phases of the cell cycle (i.e., S, G2 or M) take up more energy than the small cells in early phases (i.e., G0, G1). This is also in agreement with the observation, made with bulk measurements using cell synchronization and liquid scintillation counting, that HeLa S3 cells take up more [^18^F]FDG in both the S and G2/M phases than in G1 phase^34^.

The high [^18^F]FDG uptake by S, G2 or M cells indicates a positive correlation between FDG uptake and PI, the proportion of cells in the S, G2 or M phase. This has been confirmed in *in vivo* studies on human head and neck tumors^35,36^ and human glioma cancer^37^, but not on human lung cancer^37^. Such lack of consensus is also seen in *in vitro* studies. For example, [^18^F]FDG uptake was found to be correlated to PI in two human (SK-MEL 23 and G361) and murine (B16) melanoma cell lines, but not in SK-MEL 24 human melanoma cells^38^. Different trends were observed among three squamous-cell carcinoma cell lines; [^18^F]FDG uptake was found to be correlated with PI in UT-SCC-5 cells but not in UT-SCC-1A or UT-SCC-9 cells^39^. An inverse correlation was observed between PI and [^3^H]FDG uptake for a human ovarian adenocarcinoma cell line (HTB77IP3)^40^. Such mixed observations may be due to wide biological variations in animals and humans, particularly gene polymorphisms and environmental diversities among human populations. Single-cell radiography in tandem with various functional imaging will provide new insight into cell-level uptake of radiopharmaceuticals. This tool will help resolve confounding observations obtained with existing imaging methods and develop new radiopharmaceuticals and imaging protocols for use in clinical applications.

## Summary and Conclusion

In this paper, we have designed and built a multi-modal radiography platform that can measure the uptake of radionuclides, the cell dry mass, and the cell cycle at the single-cell level. Using this system, we have shown that HeLa cells have higher [^18^F]FDG uptake in the S, G2 or M phases than in the G1 phase, which confirms, at the single-cell, a positive correlation between [^18^F]FDG uptake and PI. We have also found a linear relationship between [^18^F]FDG uptake and cellular dry mass in HeLa cells, which suggests dry mass variations as a possible mechanism for cell cycle dependence of FDG uptake. In PET, the preferential uptake of glucose by cancerous tissues has been related to their proliferative nature, and thus the prognosis of the disease. The relationship between the two, however, has not been firmly established. Studies with this new imaging platform using various cultured and biopsied cells will provide a better understanding of the cellular mechanism that mediate FDG uptake. These findings could help improve the ability of clinicians to make accurate diagnoses and prognoses on the basis of FDG-PET scans.

## Acknowledgments

This work was supported by the University of Wisconsin-Milwaukee (startup funding to Y.S.), National Science and Engineering Research Council of Canada (to M.A.T.), and grant 5R01CA186275 of the National Institute of Health (to G.P.). The authors thank Dr. Seungeun Oh (Department of Systems Biology, Harvard Medical School) for kind donation of HeLa cells transfected with FUCCI cell cycle sensor and the team at the MGH Gordon PET Core for the production of [^18^F]FDG.

## Author Contributions

Y.S., K.T., M.N., and G.E.F. designed research; Y.S., M.A.T., and K.T. performed research; Y.S., M.A.T., K.T., J.O., G.P., M.N., and G.E.F. analyzed data; and Y.S., M.A.T., K.T., J.O., and G.P. wrote the paper.

## Additional Information

The authors declare no competing interest.

## References

1. Liberti MV, Locasale JW. The Warburg effect: how does it benefit cancer cells? Trends Biochem Sci. 2016;41(3):211–218.

2. Sengupta D, Pratx G. Imaging metabolic heterogeneity in cancer. Mol Cancer. 2016 Jan 6;15:4.

3. Pauwels EKJ, Ribeiro MJ, Stoot JHMB, McCready VR, Bourguignon M, Mazière B. FDG Accumulation and Tumor Biology. Nucl Med Biol. 1998 May 1;25(4):317–322. PMID: 9639291

4. Cherry S, Sorenson J, Phelps M. Physics in Nuclear Medicine [Internet]. Elsevier Inc.; 2012 [cited 2018 Feb 4]. Available from: https://ucdavis.pure.elsevier.com/en/publications/physics-in-nuclearmedicine-2

5. Cherry S, Sorenson J, Phelps M. Physics in Nuclear Medicine [Internet]. 2012 [cited 2018 Feb 4]. Available from: https://ucdavis.pure.elsevier.com/en/publications/physics-in-nuclear-medicine-2

6. Kinahan PE, Fletcher JW. Positron Emission Tomography-Computed Tomography Standardized Uptake Values in Clinical Practice and Assessing Response to Therapy. Semin Ultrasound CT MRI. 2010 Dec 1;31(6):496–505.

7. Veronese SM, Gambacorta M, Gottardi O, Scanzi F, Ferrari M, Lampertico P. Proliferation index as a prognostic marker in breast cancer. Cancer. 1993;71(12):3926–3931.

8. Pence JC. Prognostic Significance of the Proliferation Index in Surgically Resected Non—Small-Cell Lung Cancer. Arch Surg. 1993 Dec 1;128(12):1382.

9. Marusyk A, Polyak K. Tumor heterogeneity: causes and consequences. Biochim Biophys Acta BBA-Rev Cancer. 2010;1805(1):105–117.

10. Creath K. V phase-measurement interferometry techniques. Prog Opt. Elsevier; 1988. p. 349–393.

11. Ikeda T, Popescu G, Dasari RR, Feld MS. Hilbert phase microscopy for investigating fast dynamics in transparent systems. Opt Lett. 2005;30(10):1165–1167.

12. Iwai H, Fang-Yen C, Popescu G, Wax A, Badizadegan K, Dasari RR, Feld MS. Quantitative phase imaging using actively stabilized phase-shifting low-coherence interferometry. Opt Lett. 2004;29(20):2399–2401.

13. Liang J, Grimm B, Goelz S, Bille JF. Objective measurement of wave aberrations of the human eye with the use of a Hartmann–Shack wave-front sensor. JOSA A. 1994;11(7):1949–1957.

14. Bon P, Maucort G, Wattellier B, Monneret S. Quadriwave lateral shearing interferometry for quantitative phase microscopy of living cells. Opt Express. 2009;17(15):13080–13094.

15. Streibl N. Phase imaging by the transport equation of intensity. Opt Commun. 1984;49(1):6–10.

16. Teague MR. Deterministic phase retrieval: a Green’s function solution. JOSA. 1983;73(11):1434–1441.

17. Sung Y, Choi W, Fang-Yen C, Badizadegan K, Dasari RR, Feld MS. Optical diffraction tomography for high resolution live cell imaging. Opt Express. 2009;17(1):266–277.

18. Sung Y, Choi W, Lue N, Dasari RR, Yaqoob Z. Stain-free quantification of chromosomes in live cells using regularized tomographic phase microscopy. PloS One. 2012;7(11):e49502.

19. Pratx G, Chen K, Sun C, Axente M, Sasportas L, Carpenter C, Xing L. High-resolution radioluminescence microscopy of 18F-FDG uptake by reconstructing the β-ionization track. J Nucl Med. 2013;54(10):1841–1846.

20. Pratx G, Chen K, Sun C, Martin L, Carpenter CM, Olcott PD, Xing L. Radioluminescence microscopy: measuring the heterogeneous uptake of radiotracers in single living cells. PloS One. 2012;7(10):e46285.

21. Kim TJ, Türkcan S, Pratx G. Modular low-light microscope for imaging cellular bioluminescence and radioluminescence. Nat Protoc. 2017;12(5):1055.

22. Wang Q, Sengupta D, Kim TJ, Pratx G. Performance evaluation of 18F radioluminescence microscopy using computational simulation. Med Phys. 2017 May 1;44(5):1782–1795.

23. Edelstein A, Amodaj N, Hoover K, Vale R, Stuurman N. Computer control of microscopes using μManager. Curr Protoc Mol Biol. 2010;14–20.

24. Edelstein AD, Tsuchida MA, Amodaj N, Pinkard H, Vale RD, Stuurman N. Advanced methods of microscope control using μManager software. J Biol Methods. 2014;1(2).

25. Sobel I, Feldman G. A 3×3 isotropic gradient operator for image processing. Present Talk Stanf Art Ificial Proj. 1968;

26. Bellio R, Ventura L. An introduction to robust estimation with R functions. Proc 1st Int Work. 2005;1–57.

27. Sakaue-Sawano A, Kurokawa H, Morimura T, Hanyu A, Hama H, Osawa H, Kashiwagi S, Fukami K, Miyata T, Miyoshi H. Visualizing spatiotemporal dynamics of multicellular cell-cycle progression. Cell. 2008;132(3):487–498.

28. Bushart GB, Vetter U, Hartmann W. Glucose transport during cell cycle in IM9 lymphocytes. Horm Metab Res Horm Stoffwechselforschung Horm Metab. 1993 Apr;25(4):210–213. PMID: 8514240

29. Locasale JW, Cantley LC. Metabolic flux and the regulation of mammalian cell growth. Cell Metab. 2011;14(4):443–451.

30. Lunt SY, Vander Heiden MG. Aerobic glycolysis: meeting the metabolic requirements of cell proliferation. Annu Rev Cell Dev Biol. 2011;27:441–464.

31. Hosios AM, Hecht VC, Danai LV, Johnson MO, Rathmell JC, Steinhauser ML, Manalis SR, Vander Heiden MG. Amino acids rather than glucose account for the majority of cell mass in proliferating mammalian cells. Dev Cell. 2016;36(5):540–549.

32. Tzur A, Kafri R, LeBleu VS, Lahav G, Kirschner MW. Cell growth and size homeostasis in proliferating animal cells. Science. 2009;325(5937):167–171.

33. Sung Y, Tzur A, Oh S, Choi W, Li V, Dasari RR, Yaqoob Z, Kirschner MW. Size homeostasis in adherent cells studied by synthetic phase microscopy. Proc Natl Acad Sci. 2013;110(41):16687–16692.

34. Moriguchi H, Shozushima M. Cell cycle dependency of FDG and 67 Ga citrate uptake in HeLa S3 cells. Iwate Ika Daigaku Shigaku Zasshi. 2002;27(3):270–278.

35. Haberkorn U, Strauss LG, Reisser C, Haag D, Dimitrakopoulou A, Ziegler S, Oberdorfer F, Rudat V, van Kaick G. Glucose uptake, perfusion, and cell proliferation in head and neck tumors: Relation of positron emission tomography to flow cytometry. J Nucl Med. 1991;32(8):1548–1555.

36. Minn H, Joensuu H, Ahonen A, Klemi P. Florodeoxyglucose imaging: A method to assess the proliferative activity of human cancer in vivo. Comparison with DNA flow cytometry in head and neck tumors. Cancer. 1988;61(9):1776–1781.

37. Chung JK, Lee YJ, Kim SK, Jeong JM, Lee DS, Lee MC. Comparison of [18F] fluorodeoxyglucose uptake with glucose transporter-1 expression and proliferation rate in human glioma and non-small-cell lung cancer. Nucl Med Commun. 2004;25(1):11–17.

38. Yamada K, Brink I, Engelhardt R. Factors Influencing [F-18] 2-Fluoro-2-Deoxy-d-Glucose (F-18 FDG) Accumulation in Melanoma Cells: Is FDG a Substrate of Multidrug Resistance (MDR)? J Dermatol. 2005;32(5):335–345.

39. Minn H, Clavo AC, Grenman R, Wahl RL. In vitro comparison of cell proliferation kinetics and uptake of tritiated fluorodeoxyglucose and L-methionine in squamous-cell carcinoma of the head and neck. J Nucl Med. 1995;36(2):252–258.

40. Higashi K, Clavo AC, Wahl RL. Does FDG uptake measure proliferative activity of human cancer cells? In vitro comparison with DNA flow cytometry and tritiated thymidine uptake. J Nucl Med. 1993;34(3):414–419.

